# Bacterial Effector Screening Reveals RNF214 as a Virus Restriction Factor in Mammals

**DOI:** 10.1101/2024.11.04.621956

**Authors:** Aaron Embry, David Schad, Emily A. Rex, Neal M. Alto, Don B. Gammon

**Affiliations:** Department of Microbiology, University of Texas Southwestern Medical Center, Dallas, Texas, USA

## Abstract

Arboviruses are a group of arthropod-transmitted viruses that pose a significant threat to public health. Identifying host factors that inhibit arbovirus infection is critical for the development of strategies to prevent or treat these infections. Previously, we showed that bacterial effector proteins can be used as molecular tools to identify host immunity factors in insect cells that restrict arbovirus replication (Embry et al., 2024). Bacteria secrete effectors into the host cell cytoplasm to inhibit various innate immune defenses. Here, we apply our bacterial effector screening system to identify host antiviral immunity factors in two mammalian hosts – bats and humans. By screening a library of 210 effectors encoded by seven distinct bacterial pathogens, we identified three bacterial effectors (IpaH4, SopB, and SidM) that enhance the replication of both togaviruses and rhabdoviruses in bat and human cells. We also discovered several effectors that enhance arbovirus replication in a virus- or host-specific manner. We further characterize the mechanism by which the *Shigella flexneri* encoded E3 ubiquitin ligase, IpaH4, enhances arbovirus infection in mammalian cells. Using yeast two-hybrid, ubiquitin-activated interaction traps, *in vitro* ubiquitination assays and cellular approaches, we show the uncharacterized mammalian RING-domain containing protein, RNF214, to be directly targeted by IpaH4 for ubiquitination-mediated degradation. Phylogenetic analyses of RNF214 proteins indicate they are widely conserved among many vertebrate species, suggesting an important evolutionary function. We show that RNF214 overexpression suppresses arbovirus infections in a manner dependent upon its putative E3 ubiquitin ligase activity, while RNF214 depletion enhances these infections in human and bat cells. These data suggest that RNF214 proteins are important innate immune factors involved in combating viral infection. Collectively, our work shows that bacterial effectors can be useful tools for uncovering novel mammalian antiviral machinery.

**Author Summary:** Arboviruses are viruses transmitted by arthropod vectors such as mosquitoes or biting flies to both animal and human hosts. Arboviruses cause diseases ranging from mild febrile illnesses to fatal encephalitic infection. Here, we use bacterial effector proteins to identify mammalian factors that block arbovirus replication. Bacterial effectors encoded by pathogenic bacteria are secreted into mammalian host cells to inhibit cellular antimicrobial responses. Like viruses, many pathogenic bacteria replicate inside of mammalian host cells, thus we hypothesized that some bacterial effectors may target and inhibit the host immune response that are normally restrictive to both bacteria and viruses. After screening a library of >200 bacterial effector proteins, we identified three effectors that promote the replication of arboviruses belonging to two distinct families in bat and human cells. We further show that one of our most potent effectors, IpaH4, enhances arbovirus replication by targeting mammalian RNF214 proteins for degradation. RNF214 proteins are poorly characterized, but our phylogenetic analyses suggest these proteins are widely conserved among vertebrate organisms. We show that depletion of RNF214 protein levels in either bat or human cells sensitizes these cells to arbovirus infections, revealing a new role for RNF214 proteins in antiviral defense. Our study demonstrates the utility of bacterial effectors as tools for identifying new host immune machinery in mammalian hosts.

## Introduction

Arboviruses are transmitted by arthropod vectors to a wide variety of mammalian hosts. Whether arbovirus infection ensues after vector transmission depends upon the ability of arboviruses to overcome mammalian antiviral defenses. Bats are diverse group of mammals encompassing >1000 species and are found in all global regions inhabited by humans. Importantly, bats act as natural reservoirs for viruses affecting humans, including highly pathogenic coronaviruses and filoviruses [1–4]. Moreover, accumulating evidence suggests that bats may also be reservoirs for arboviruses that infect humans, such as togaviruses [e.g. Sindbis virus (SINV), Ross River virus (RRV), and O’nyong’nyong virus (ONNV)] [5–8]. However, little is known regarding the bat antiviral responses that combat arboviruses. Due to a lack of screening tools (e.g. CRISPR-Cas9 libraries), our understanding of bat immune systems is extremely limited and has largely been gleaned from comparative genomics and transcriptomic profiling of virus-infected bat cells [9–13]. Therefore, developing functional assays to identify bat factors relevant to arbovirus restriction is paramount.

We recently identified host antiviral factors in invertebrate cells by screening for bacterial effectors that enhance arbovirus replication in normally restrictive lepidopteran insect cells [14]. Bacterial effectors are proteins injected into eukaryotic host cells through needle-like secretion systems to subvert the immune response. Therefore, if bacterial effectors inhibit host immune factors that restrict both bacteria and viruses, they may enhance virus replication when expressed in restrictive cells. Moreover, these effectors can be used as molecular tools to identify the host immune machinery they target [14].

Here we show that bacterial effectors can be used as functional tools to identify arbovirus restriction factors encoded by mammalian species such as bats and humans. By screening a library of 210 bacterial effectors, we identify dozens of effectors that rescue either togavirus or rhabdovirus infection in either humans or bats, suggesting some of these target virus or host-specific restrictions. Moreover, we discovered three effector proteins that enhance both togavirus and rhabdovirus replication in both host species. Characterization of one of the most potent effector proteins, *Shigella flexneri*-encoded IpaH4, led us to identify mammalian RNF214 proteins as direct targets of IpaH4 and novel virus restriction factors in bat and human cells. Our work demonstrates that bacterial effector screening provides a functional platform for uncovering effector functions and mammalian antiviral machinery.

## Results

### Inhibition of Host Transcription Enhances Arbovirus Replication in R06E Bat Cells

Previously, we showed that the togavirus RRV and the rhabdovirus vesicular stomatitis virus (VSV) undergoes an abortive infection in *Lymantria dispar*-derived LD652 cells, but were rescued by actinomycin D (ActD) treatment, which globally inhibits cellular transcription [14, 15]. This suggested that viral replication could be enhanced in insect cells when host responses were inhibited, giving us confidence to screen for bacterial effectors that similarly enhanced arbovirus replication in these cells [14]. Thus, we asked if ActD treatment also enhances arbovirus replication in bat cells in order to justify a similar bacterial effector screen in mammalian cells.

We examined arbovirus replication in *Rousettus aegyptiacus* (Egyptian fruit bat)-derived R06E cells for several reasons: 1) the *R. aegyptiacus* genome has been sequenced [9], which is critical for identifying host antiviral factors encoded by this species; 2) R06E cells are capable of supporting arboviruses replication; 3) evidence suggests that *R. aegyptiacus* animals are naturally infected with viruses belonging to *Togaviridae* and *Rhabdoviridae*, the two viral families we focus on in this study [5, 6]; and 4) *R. aegyptiacus* is a reservoir for other pathogenic viruses such as Marburg virus [2]. Thus, an improved understanding of *R. aegyptiacus* immunity may have implications for other viral diseases, beyond arboviruses.

To determine if ActD treatment alters arbovirus infection of R06E cells, we challenged cells with GFP reporter arboviruses (RRV-GFP [16] and VSV^M51R^-GFP [15]) across increasing ActD concentrations. It is important to note that VSV^M51R^-GFP is an attenuated strain with a reduced ability to block antiviral host responses [17–20], and thus we hypothesized that it would be easier to detect enhanced replication with this mutant. Interestingly, ActD increased GFP signals during RRV-GFP or VSV^M51R^-GFP infection in a dose-dependent manner, peaking at ∼60-to 70-fold at the highest dose tested (10 nM) without significantly impacting cell viability via a lactate dehydrogenase (LDH)-based cytotoxicity assay (**Fig S1A-C**). These results suggest that R06E cells express antiviral factors that restrict arbovirus replication.

### Specific Bacterial Effectors Enhance Arbovirus Replication in Bat and Human Cells

Expression of specific bacterial effectors in lepidopteran cells can rescue otherwise abortive arbovirus infections [14]. To determine if effectors could enhance arbovirus replication in mammalian cells, we adapted our library of 210 effector genes [14] for lentivirus-based expression [21]. To compare potential arbovirus-enhancing activities of effectors in additional mammalian species, we conducted effector screens in both bat R06E cells and human U2OS cells, the latter being a human cell type known to be susceptible to RRV and VSV [22, 23].

R06E and U2OS cells were transduced with the effector library for 48 h and then infected with either RRV-GFP or VSV^M51R^ -GFP for 20 h. Cells were then stained with CellTracker Blue dye (to normalize viral GFP signals to cell number), fixed, then subjected to automated fluorescence microscopy [14]. Effector proteins that enhanced viral GFP signals by >10-fold over cells transduced with lentiviruses expressing firefly luciferase (control) were considered “hits”, a cut-off we established to avoid examining false-positives (**Fig 1AB**). Importantly, we assessed transduced cells for cytotoxicity due to effector expression and eliminated 18 effectors in bat and 4 in human cells from further consideration due to toxicity (**Fig S1DE**). We identified 15 unique enhancers of RRV-GFP infection in R06E cells and 23 in U2OS cells. Furthermore, 18 effectors enhanced VSV^M51R^-GFP in R06E and 11 enhanced infection of U2OS cells. We found two effectors to enhance both togavirus and rhabdovirus replication in both bat and human cells: SopB and IpaH4 (**Fig 1AB** and **Fig S2AB**). While SidM failed to enhance RRV-GFP in U2OS cells, it was a hit in three other screens (**Fig S2AB**). Thus, we decided to focus our downstream analyses on these three effectors given their broad rescue phenotypes. These effectors are encoded by three distinct bacterial pathogens: *Salmonella enterica* serovar Typhimurium (SopB), *Shigella flexneri* (IpaH4), and *Legionella pneumophilia* (SidM). Collectively, these results suggest that specific effectors encoded by diverse bacterial pathogens can enhance togavirus and rhabdovirus replication in mammalian cells.

**Fig 1.**
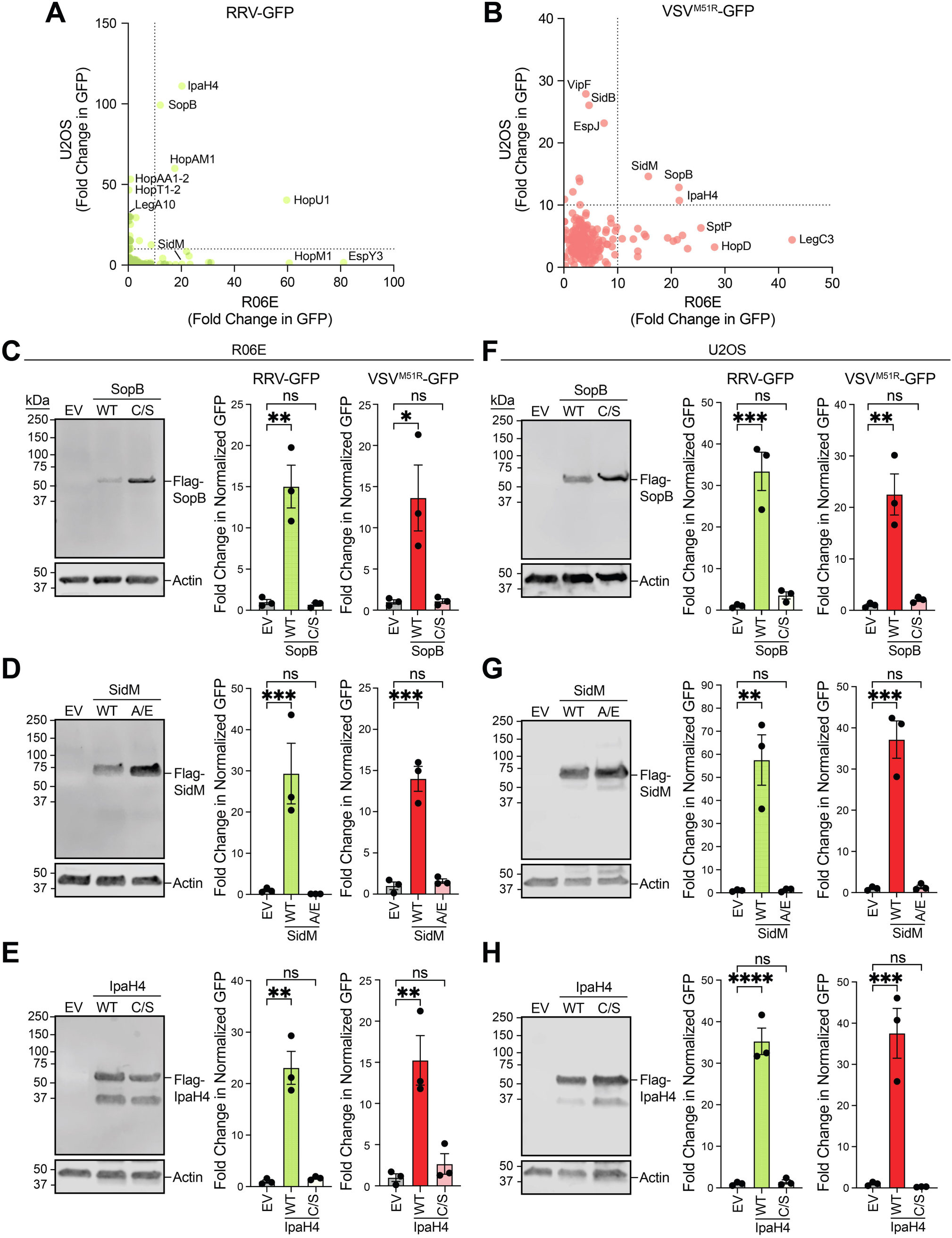
Identification of bacterial effectors that promote arbovirus replication in bat and human cells. **A-B**. Fold-change in viral GFP signals (normalized to CellTracker Blue signal) relative to cells transduced with lentivirus expressing firefly luciferase (LUC) (negative control) for RRV-GFP (A) and VSV^M51R^-GFP (B) screens 20 hpi. The cutoff for effectors to be scored as hits was a >10-fold change in over negative controls (cutoffs represented by dotted lines). Data points are means. **C-E.** Representative immunoblots of Flag-tagged bacterial effector expression in bat R06E cells 48 h post-transfection with pCDNA3.1 vectors. Additionally, fold-change in normalized viral GFP signal relative to cells transfected with empty vector (EV) is shown. Cells were stained 20 hpi with CellTracker Orange dye and imaged to calculate fold-change in normalized GFP signal over signals in EV treatments. Wild-type (WT) effectors are compared to their mutants (SopB^C420S^, SidM^A435E^, IpaH4^C339S^). **F-H.** Representative immunoblots and fold-change in normalized viral GFP signal for effectors experiments conducted as in C-E but in human U2OS cells. Quantitative data in C-H are means ± SEM; n=3. Statistical significance in graphs in C-H was determined with One-way ANOVA; ns (not significant), *=P<0.05, **=*P*<0.01, ***=*P*<0.001, ****=*P*<0.0001.

### Validation of Effector Activities Required for Viral Enhancement

Because the effector genes in our expression library are not epitope-tagged, we wanted to confirm the expression and rescue functions of key hits from our screens. To do this, we cloned Flag-tagged SopB, IpaH4, and SidM into pcDNA3.1 and transfected cells with these vectors for 48 h to confirm expression. SopB is an inositol-phosphatase [24], IpaH4 is an E3 ubiquitin ligase [14], and SidM is a guanine nucleotide exchange factor for Rab1 [25, 26]. Therefore, we generated point mutant versions of these effectors that inactivate these known catalytic activities or functions to determine if these activities were important for promoting arbovirus replication. While both wild-type (WT) and mutant forms of these effectors were detected by immunoblot, only WT effectors significantly enhanced arbovirus infection (**Fig 1C-H**), suggesting these diverse effector functions are required to relieve arbovirus restrictions in mammalian cells.

### IpaH4 E3 ubiquitin ligase activity is required for ubiquitination and degradation of host RNF214

Our previous study demonstrated that the *Shigella flexneri*-encoded IpaH4 is a bacterial effector with E3 ubiquitin ligase activity [14]. We identified putative human binding partners using a yeast two-hybrid (Y2H) approach with IpaH4 bait. One of the top hits was the RING-finger domain (RNF) containing protein RNF214, a putative E3 ubiquitin ligase. Nine independent prey clones targeting a central region of RNF214 spanning amino acid (a.a) 276-504 were identified (**Fig 2A**) [14]. Interestingly RNF214 was the only human protein identified as a potential substrate for IpaH4 using ubiquitin-activated interaction trap assays [27] that overlapped with our Y2H hits [14]. We did not pursue RNF214 in our previous study examining IpaH4 targets in *L. dispar* cells, as RNF214 proteins are absent in moths and appear to be largely restricted to vertebrates (**Fig 2B-D**). For example, through reciprocal BLAST and phylogenetic analyses, we identified RNF214 orthologs in numerous mammalian species. However, RNF214 orthologs were rarely found in invertebrate species other than the notable exceptions of starfish and tubeworms (**Fig 2B-D**). Interestingly, AlphaFoldv2 predictions suggest human RNF214 is largely disordered, except for α-helices encompassing most of the region predicted to interact with IpaH4 by Y2H (a.a. 216-422), and portions of the C-terminal RING domain (a.a 550-688) (**Fig 2E**).

**Fig 2.**
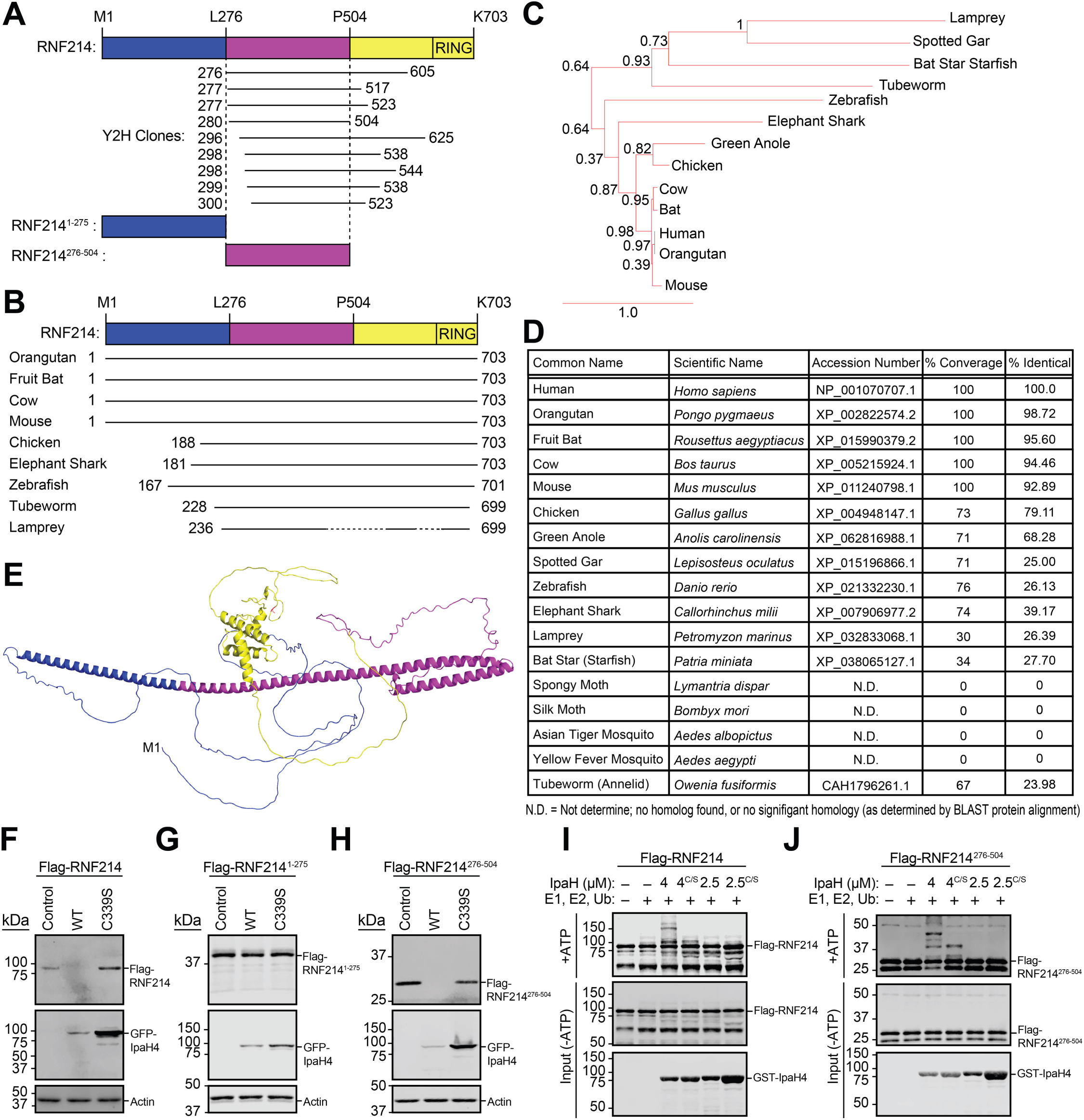
The central domain of the conserved host protein, RNF214, is sufficient for IpaH4-mediated ubiquitination and degradation. **A.** Schematic of human RNF214 with mapping of prey clones collectively identified in two yeast two-hybrid screens using IpaH4 bait [14]. Numbers indicate RNF214 a.a. encoded by each prey clone. **B.** Schematic displaying the relative conservation and length of RNF214 across indicated organisms. **C.** PhyloTv2-generated phylogenetic tree of RNF214 orthologs. **D.** Table of RNF214 orthologs in various vertebrate and invertebrate species generated via reciprocal BLAST analysis. **E.** AlphaFold2-predicted structure of human RNF214. **F.** Representative immunoblot of degradation assay using Flag-tagged RNF214 co-transfected into U2OS cells with GFP-IpaH4 (WT) or GFP-IpaH4^C339S^ (C339S) catalytic mutant expression vectors. **G-H.** Representative immunoblot of degradation assays using Flag-tagged RNF214 truncations, RNF214^1-275^ (**G**) or RNF214^276-504^ (**H**), co-transfected into U2OS cells with GFP-IpaH4 (WT) or GFP-IpaH4^C339S^ (C339S) vectors. **I-J.** Representative immunoblot of *in vitro* ubiquitination assay showing direct IpaH4-mediated ubiquitination of purified, recombinant human Flag-RNF214 (**I**) and Flag-RNF214^276-504^ (**J**) proteins.

To determine if IpaH4 targets RNF214, we conducted degradation assays [14] in U2OS cells and found Flag-RNF214 levels to decrease in the presence of WT, but not the catalytically inactive (IpaH4^C339S^), IpaH4 protein (**Fig 2F**). Consistent with our Y2H results, expression of the central RNF214 region (a.a. 276-504) also led to its degradation in the presence of WT IpaH4 (**Fig 2G**). However, IpaH4 did not alter the levels of a N-terminal fragment of RNF214 (a.a. 1-275) (**Fig 2H**). This suggests that the central domain of RNF214 (a.a. 276-504) is sufficient for IpaH4-mediated degradation. *In vitro* ubiquitination confirmed that purified, recombinant Flag-RNF214 and Flag-RNF214^276-504^ are targeted for ubiquitination by GST-tagged WT IpaH4, but not the IpaH4^C339S^ catalytic mutant [14] (**Fig 2IJ**). Moreover, *S. flexneri* IpaH2.5, a related E3 ubiquitin ligase [28], was unable to ubiquitinate RNF214, suggesting it is a specific target of IpaH4 (**Fig 2IJ**). These results indicate that IpaH4 directly ubiquitinates the central domain of RNF214 for degradation in mammalian cells.

### RNF214 overexpression suppresses arbovirus infection of mammalian cells

To determine if RNF214 was relevant to arbovirus restriction, we overexpressed Flag-RNF214 proteins in bat and human cells for 48 h and then challenged cells with arboviruses. To test if the putative E3 ubiquitin ligase activity was important for viral restriction, we used bioinformatics to align the RNF domain of RNF214 with other characterized E3 ligases to identify catalytic residues (**Fig 3A**). RNF domains consist of 40-80 a.a. residues, within which a combination of eight cysteine and histidine residues coordinate two zinc ions to create a distinct three-dimensional structure [29]. Our alignment identified RNF214 a.a. C658 as a likely catalytic residue that was also conserved between bat and human proteins (**Fig 3B**). Overexpression of WT Flag-RNF214 significantly inhibited RRV-GFP and VSV^M51R^-GFP in bat and human cells. However, Flag-RNF214^C658S^ overexpression did not alter arbovirus replication in bat cells or affect VSV^M51R^-GFP replication in human cells, despite expressing to similar levels as WT Flag-RNF214. However, there was minor, but statistically significant suppression of RRV-GFP by Flag-RNF214^C658S^ in U2OS cells (**Fig 3C-F**). These data suggest RNF214 suppresses arbovirus replication in bat and human cells via a mechanism largely dependent on its putative E3 ubiquitin ligase activity.

**Fig 3.**
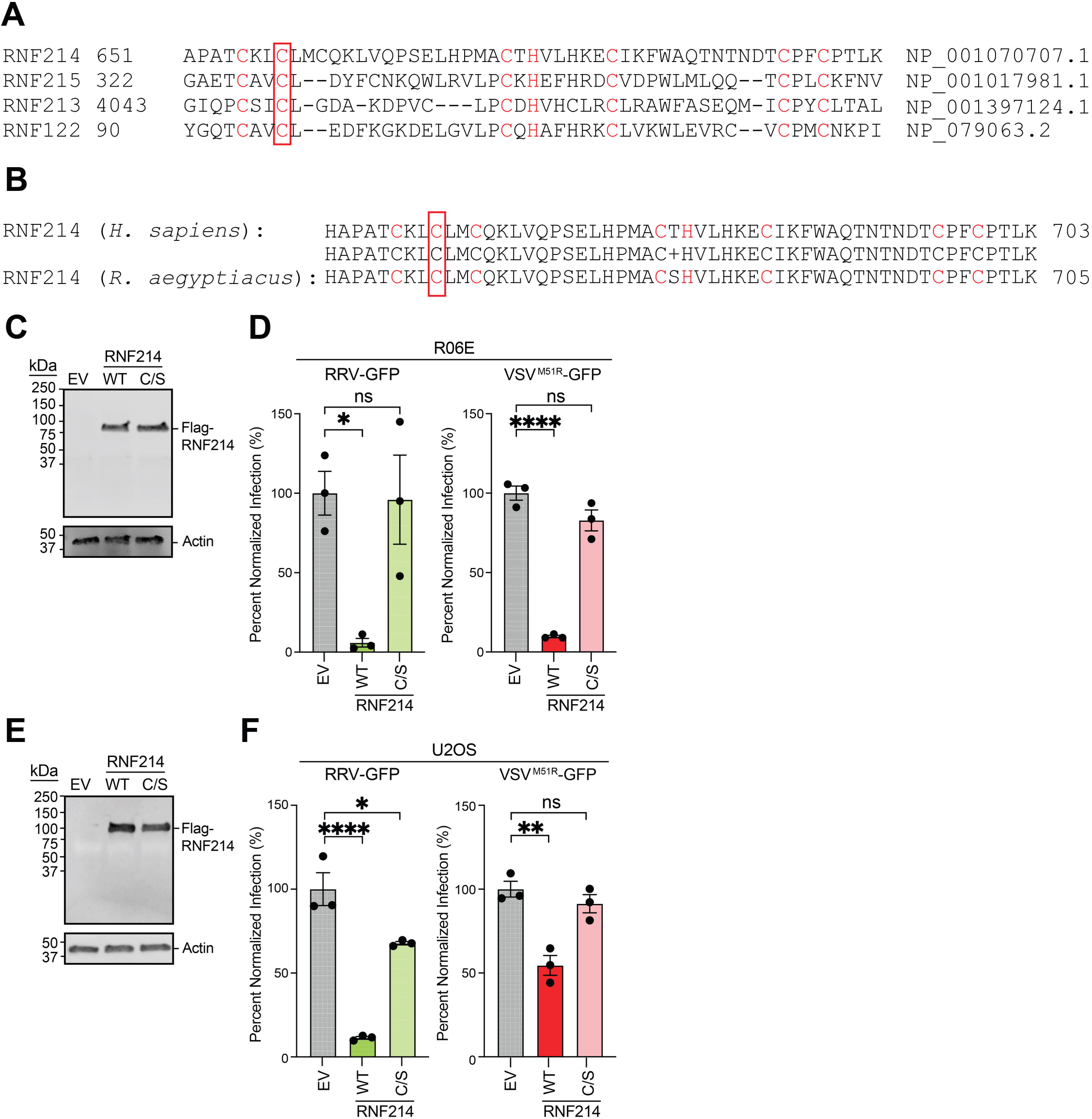
Overexpression of RNF214 suppresses arbovirus replication in bat and human cells. **A.** a.a. alignment of human RNF214 with other human RNF containing proteins. Residues in the characteristic RNF domain “C4-H-C3” motif are highlighted in red, and the C658 residue targeted for substitution mutation is boxed. **B.** a.a. alignment of human RNF214 with *R. aegyptiacus* RNF214. Residues are highlighted as described in A. **C.** Representative immunoblot of Flag-tagged RNF214 and RNF214^C658S^ (C/S) overexpression in R06E cells for 48 h. **D.** Percent of GFP signal relative to empty vector (EV) control after R06E cells overexpressing RNF214 constructs (WT or RNF214^C658S^) were infected with RRV-GFP (MOI=0.05) or VSV^M51R^-GFP (MOI=0.005) were for 20 h. **E.** Representative immunoblot of Flag-tagged RNF214 constructs overexpressed in U2OS cells for 48 h. **F.** Percent of GFP signal relative to empty vector (EV) control in R06E cells overexpressing Flag-RNF214 constructs (WT or RNF214^C658S^) RRV-GFP (MOI=0.05) or VSV^M51R^-GFP (MOI=0.005) for 20 h. Data are means ± SEM; n=3. Statistical significance was determined with One-way ANOVA; ns (not significant), *=P<0.05, **=*P*<0.01, ***=*P*<0.001, ****=*P*<0.0001.

### RNF214 Depletion Enhances Virus Replication in Mammalian Cells

To confirm a role for RNF214 in virus restriction, we knocked down RNF214 in R06E and U2OS cells using RNAi and then challenged cells with arbovirus infection. RRV-GFP and VSV^M51R^-GFP infection was enhanced in two of three RNF214 siRNA treatments (**Fig 4AB**). Furthermore, RNF214 depletion in these cells enhanced the replication of another GFP encoding togavirus, ONNV-GFP (**Fig 4C**) [14]. We confirmed the specificity of RNF214 RNAi in R06E cells (**Fig 4D**). RNF214 knockdown also enhanced RRV-GFP and VSV^M51R^-GFP in U2OS cells (**Fig S3A-C**). To further confirm these results, we used CRISPR-Cas9 genomic editing to knockout RNF214 in U2OS cells (U2OS^ΔRNF214^). Immunoblotting confirmed RNF214 knockout (**Fig 4E**). Infection of U2OS^ΔRNF214^ cells with RRV-GFP, VSV^M51R^-GFP, ONNV-GFP, and SINV-GFP [14] resulted in 10-1000-fold increases in GFP signal compared to infections in control U2OS cells. Supernatants from U2OS^ΔRNF214^ cells also exhibited higher viral titers (**Fig 4F-I**).

**Fig 4.**
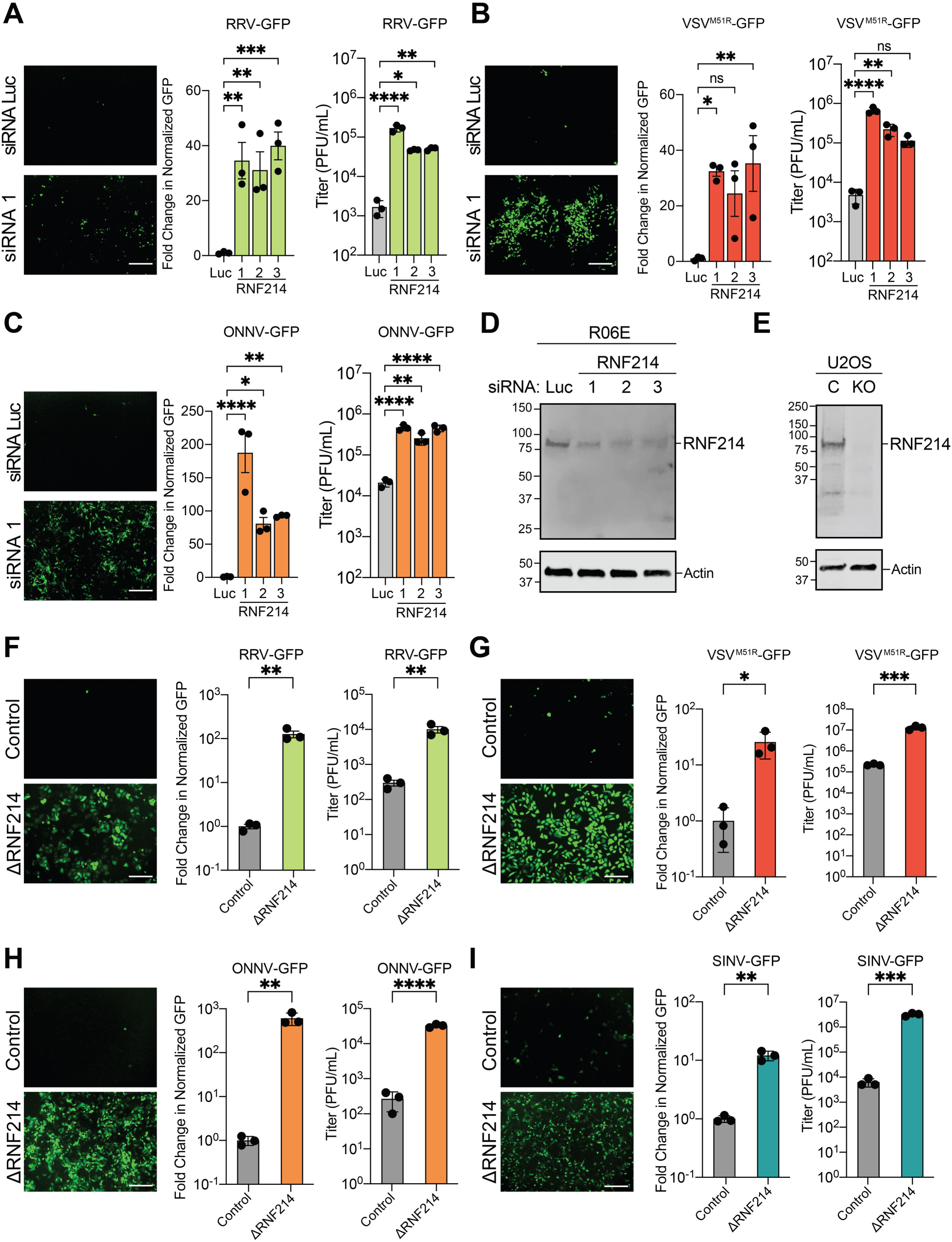
Depletion of RNF214 enhances arbovirus replication in bat and human cells. **A.** Representative fluorescence microscopy images (GFP channel) of R06E cells transfected with firefly luciferase (Luc)- or RNF214-targeting siRNAs. 48 h post-knockdown, cells were infected with RRV-GFP (MOI=0.001) for 20 h. Cells were then stained with CellTracker Orange, imaged, and fold-change in GFP signal compared to the Luc control was calculated. Supernatants were also collected to assess viral titers by plaque assay. **B-C.** Similar experiments were performed as in A but for VSV^M51R^-GFP (MOI=0.0001) (**B**) and ONNV-GFP (MOI=0.001) (**C**). **D.** Representative immunoblot of endogenous bat RNF214 levels following siRNA knockdown in R06E cells for 48 h. **E.** Representative immunoblot of endogenous human RNF214 levels following in U2OS cells transduced with control lentivirus (C) or lentivirus expressing guide RNAs targeting RNF214 (KO). **F.** Representative fluorescence microscopy images (GFP channel) in control or ΔRNF214 U2OS cells infected with RRV-GFP (MOI=0.001) 20 h. Cells were then stained with CellTracker Orange, imaged, and fold-change in GFP signal compared to Control U2OS cell infections was calculated. Supernatants were also collected to assess viral titers by plaque assay. **G-I.** Similar experiments were performed as in F but for VSV^M51R^-GFP (MOI=0.0001) (**G**), ONNV-GFP (MOI=0.001) and SINV-GFP (MOI=0.001) (**H**). Data are means ± SEM; n=3. Statistical significance for A-C was determined with One-way ANOVA and with unpaired student’s t-test for F-I; ns (not significant), *=P<0.05, **=*P*<0.01, ***=*P*<0.001, ****=*P*<0.0001. Scalebars indicate 200 μm on all microscopy images.

To assess if other viruses are enhanced in U2OS^ΔRNF214^ cells, we challenged these cells with dsDNA viruses, vaccinia virus (VACV-FL-GFP) [30] and herpes simplex virus 1 (HSV-1-GFP) [31] but observed no significant changes in GFP signal or viral titer (**Fig S4AB**). In contrast, wild-type VSV-GFP was moderately enhanced in U2OS^ΔRNF214^ cells (**Fig S4C**), suggesting that even the non-attenuated VSV strain can be restricted by RNF214. In contrast, other negative-sense ssRNA viruses, such as the paramyxoviruses, human parainfluenza virus 1 (HPIV1-GFP) [32] and Sendai virus (SeV-GFP) [31] either exhibited no change or minor increases in viral replication (**Fig S4DE**). A similar minor enhancement was observed in U2OS^ΔRNF214^ cells for the positive-sense ssRNA flavivirus, West Nile virus (WNV-GFP) [21] (**Fig S4F**). Interestingly, Venezuelan equine encephalitis virus (VEEV-GFP) [33, 34] showed a ∼15-fold increase in titer in U2OS^ΔRNF214^ cultures (**Fig S4G**), consistent with our observations for other togaviruses (**Fig 4F**). In summary, these data suggest RNF214 is important for restricting ssRNA virus infections but not DNA viruses (**Fig 4H**).

## Discussion

Here, we applied our bacterial effector screening methodology to identify immune evasion proteins that modulate antiviral immunity in mammals. Such functional screens are important to develop in non-model species, such as bats, given the lack of tools to perform more classical screens (e.g. CRISPR-Cas9) for host immunity factors on a high throughput scale. By applying our screening approach to both bat and human cells, we identified 30 effectors that enhance arbovirus infection in bat cells and 29 in human cells. Moreover, we discovered three effectors: SopB, SidM, and IpaH4, that enhanced togavirus and rhabdovirus replication in bat and human cells.

Here we focused on understanding how the *S. flexneri*-encoded E3 ubiquitin ligase, IpaH4, enhances arbovirus replication in mammalian cells. Our Y2H [14] and ubiquitin-activated interaction trap assays [27] with IpaH4 identified human RNF214 as a putative substrate, but it was unclear if RNF214 was relevant to innate immunity. This was important to determine as our phylogenetic analyses suggest RNF214 proteins are widely conserved among vertebrate species including mammals. We found IpaH4 to directly ubiquitinate RNF214 *in vitro* and promote its degradation, indicating it is a bona fide substrate. Furthermore, RNF214 overexpression inhibited, and RNF214 depletion enhanced, arbovirus infection of bat and human cells, suggesting RNF214 proteins are potent arbovirus restriction factors in mammalian hosts. Moreover, by screening a panel of 11 viruses, representing 6 viral families, we found RNF214 to most potently inhibit togaviruses and rhabdoviruses, but also restricts specific paramyxoviruses and flaviviruses. These data strongly implicate mammalian RNF214 proteins in antiviral immunity. Interestingly, other RNF domain-containing proteins, such as RNF213, have been shown to restrict both viruses and bacteria [35–37]. Therefore, RNF214 may be an additional member of an ever-expanding group of host E3 ubiquitin ligases that use ubiquitination to combat microbial infection [38–40]. Consistent with this, the RNF214^C658S^ mutant, predicted to be catalytically inactive, was unable to suppress arbovirus replication in mammalian cells.

Future studies will aim to confirm RNF214 E3 ubiquitin ligase activity *in vitro* and determine the mechanism by which these host factors restrict viral replication. Interestingly, two recent studies suggest that RNF214 promotes ubiquitination of TEAD proteins to regulate the Hippo pathway, which plays roles in cellular homeostasis and proliferation [41]. Several viruses suppress this pathway, including SARS-CoV-2 and Ebola virus, suggesting it may play a role in antiviral defense [42, 43]. Whether RNF214-mediated regulation of the Hippo pathway explains its antiviral activity will be important to determine.

In summary, our study illustrates how bacterial effector screens can be applied to mammalian systems to uncover new effector protein functions and novel innate immunity factors. Thus, our system provides a power approach to reveal critical pathogen-host interactions that regulate host susceptibility to infection.

## Materials and Methods

### Cell Lines and Cell Culture

Mammalian cell lines were maintained at 37°C in 5% CO_2_ atmosphere. U2OS cells were cultured in DMEM supplemented with 10% FBS containing 1% non-essential amino acids (NEAA), 1% L-glutamine, and 1% antibiotic/antimycotic (Gibco) (**Table S2**). R06E cells were cultured in DMEM:F12 supplemented with 10% FBS and 1% antibiotic/antimycotic (Gibco). BSC-40 cells were cultured in MEM supplemented with 5% FBS (**Table S2**).

### Viruses

Stock preparation, culture of recombinant viruses, and titration by fluorescent foci/plaque assay on BSC-40 cells was performed as previously described [14, 15]. Viral innocula were incubated with cells for 1 h in serum free media before the addition of complete media for the remainder of the infection. Where indicated, complete media containing ActD at the indicated dose was added for the remainder of the infection.

### Cell Viability Assay

Cell viability was measured using a CyQUANT^TM^ LDH Cytotoxicity assay (Invitrogen; **Table S2**) as previous described [14] 48 h post-transduction of cells with the effector expression library.

### Plasmid Constructs for Mammalian Cell Expression

The 210 bacterial effector protein coding sequences was generated in pENTR/D as previously described [44]. Briefly, PCR-amplified bacterial genomic DNA sequences were cloned into pENTR/D-TOPO (Invitrogen) using topoisomerase I. Source organisms were *Pseudomonas syringe* pv. Tomato (ATCC BAA-871D-5), *Legionella pneumophila* Philadelphia-1 (ATCC 33152D-5), *Salmonella Typhimurium* LT2 (gift of Jack Dixon, University of California San Diego), EHEC H7:O157 (gift of Vanessa Sperandio, University of Wisconsin-Madison), *Shigella flexneri* M90T (gift of Jack Dixon, University of California San Diego), and *Bartonella henselae* Houston-1 (gift of Alexei Savchenko, University of Toronto). For expression in mammalian cells, the library was cloned into the lentiviral expression vector pTRIP-CMV-IVSb-IRES-TagRFP [21] using Gateway LR Clonase II (Invitrogen) (**Table S2**).

RNF214 was Gibson cloned from pENTR221 (DNASU HsCD00513497) into pFLAG-CMV-6b, imparting an N-terminal Flag tag. Flag-tagged RNF214 fragments were also generated in pFLAG-CMV-6b by Gibson cloning. For bacterial expression, sequence encoding Flag-RNF214 or only the 276-504 a.a. RNF214 fragment were Gibson cloned into pProEX-HTb (Invitrogen), imparting a 6xHis tag at the N-terminus upstream of the Flag sequence.

N-terminal Flag-tagged versions of SopB (WFG56166.1), SidM (YP_096471.1), and C-terminal Flag-tagged IpaH4 (EID62426.1) wild-type and point mutants were generated as previously described and cloned into pcDNA3.1 for expression in mammalian cells [14].

### General Transfection Protocols

Unless otherwise stated, 100,000 cells were plated into 24-well dishes, transfected with 500 ng of expression vectors for 48 h prior to manipulation (e.g. protein extraction or virus infection). For U2OS cell expression of the pcDNA3.1 constructs, 500 ng of vector was mixed with 100 μL of OptiMEM media and 1.5 μL of Lipofectamine 2000 Reagent (**Table S2**). This mixture was incubated at room temperature for 20 min, and media on the cells was changed to 500μL OptiMEM, then the transfection mixture was added dropwise to wells, incubated overnight and media was then changed to complete DMEM ∼16 h post-transfection.

For R06E transfection experiments, as well as HEK293T transfection, 500 ng of vector was mixed with 50 uL of Opti-MEM and 1 μL of FuGENE HD Transfection Reagent (Promega; **Table S2**). This mixture was incubated at room temperature for 20 min before adding dropwise to the well. Cells were then incubated for 48 h prior to further manipulation or infection.

Transient siRNA-mediated knockdown was achieved by reverse transfection of R06E cells with 8 pmol siRNA and 1.5 μL of FuGENE HD Reagent according to the manufactureer’s protocol. The same protocol was used for U2OS cells except Lipofectamine 2000 was used as the transfection reagent according to the manufacturer’s protocol. Cells were transfected for 48 h and then either subjected to protein extraction and immunoblotting or were subjected to GFP reporter virus infection to assess replication phenotypes.

### Immunoblotting

Immunoblotting was conducted as previously described with primary antibodies, secondary antibodies conjugated with infrared dyes and a Li-Cor Odyssey scanner [14].

### Bacterial Effector Screens and Fluorescence Microscopy

Cells were seeded into 96-well clear bottom dishes, transduced with the lentivirus effector library in quadruplicate for 48h, and then challenged with indicated GFP reporter viruses. At the indicated times post-infection, cells were stained as previously described [14] with CellTracker Dye, fixed with PFA, and cells were imaged by the UT Southwestern Medical Center High-Throughput Screening Core using an IN Cell Analyzer 6000 (Molecular Devices) scope equipped with 405, 488, and 561 nm lasers using a using a 10x objective. Four images were taken/well so that fluorescence signals/well resulted from an average of these four images. Signals across all 4 replicate wells for each effector treatment were then used to determine GFP reporter virus infection levels. Image analysis was conducted using Fiji 2.14.0 (NIH) to quantify the percent area of each field of view containing GFP signal and these signals were normalized to CellTracker Dye signal to account for cell number [14]. Finally, normalized GFP signals for each effector treatment were plotted as a fold change in GFP signal relative to cells transduced with lentivirus vector expressing firefly luciferase (LUC) (negative control). Validation assays to confirm virus-enhancing phenotypes after transfection of pCDNA3.1 expression plasmids encoding Flag-tagged effector constructs were conducted and analyzed in a similar manner except empty pCDNA3.1 vector was used as a negative control treatment.

### RNF214 Degradation Assay

∼75,000 cells were co-transfected with 150 ng target pCDNA3.1 vectors encoding full-length Flag-RNF214 (or RNF214 fragments) and 350 ng of pEGFP-C2 encoding either GFP, GFP-IpaH4, or GFP-IpaH4^C339S^[14] using 1.5 μL of X-tremeGENE 9 and 50 μL of OptiMEM (Sigma; Table S2). After 24 h, cells were harvested directly in 1X Laemmeli buffer containing β-mercaptoethanol. Protein extracts were then subjected to SDS-PAGE and subsequent immunoblotting with indicated antibodies.

### Protein Purification

For human Flag-RNF214 full length or fragment expression *E. coli* BL21 cells were transformed with pProEX-HTb-6xHis-FLAG-RNF214 or pProEX-HTb-6xHis-FLAG-RNF214^276-504^ vectors and spread onto LB agar plates containing 100 µg/mL ampicillin. A colony was picked and grown in a 30 mL LB culture with 100 µg/mL ampicillin overnight at 37°C with shaking. The following day, this culture was added to 1 L of LB with ampicillin and grown at 37°C with shaking to an OD600 of 0.8. Protein expression was then induced by the addition of 0.5 mM IPTG and the culture was grown overnight at 18°C with shaking. Bacteria were pelleted at 4000xg for 20 min (Beckman Avanti J-25) and resuspended in 30 mL of purification buffer (20 mM HEPES, 150 mM NaCl, 1 mM TCEP, and 1X SigmaFAST protease inhibitor (Sigma S8820), pH=7.5). Lysate was clarified by centrifugation at 10,000xg for 15 min. The fusion protein was then affinity-purified over an Econo-Pac chromatography column (Biorad 9704652) using TALON Metal Affinity Resin (Takara 635502) and eluted in purification buffer containing 500 mM imidazole. The protein was dialyzed using a 10,000 MWCO Slide-A-Lyzer dialysis cassette (Thermo 66810) in 1 L of purification buffer without imidazole. Protein was combined with glycerol at 20% and stored at -80°C. Recombinant GST-IpaH4 for *in vitro* ubiquitination experiments was expressed and purified as previously described [14].

### In-Vitro Ubiquitination Assay

*In vitro* ubiquitination reactions were performed as previously described [14] with minor modifications. Briefly, reactions were performed in 50 mM HEPES pH 7.5, 150 mM NaCl, 20 mM MgCl_2_, and 10 mM ATP in a volume of 30 µL. The following recombinant proteins were added: 1 µM UbE1 (E1), 5 µM UbcH5b (E2), 5 µM GST-IpaH or GST-IpaH4^C339S^, 50 µM ubiquitin, and 5 µM 6xHis-FLAG-tagged RNF214 or its truncated versions. ATP was added last to initiate reactions, which were then incubated for 2 h at 30°C before the addition of 30 µL 2X Laemmli buffer containing β-mercaptoethanol. Samples were then boiled for 10 min at 95°C, subjected to SDS-PAGE, and immunoblotting.

### Lentivirus Production

HEK293T cells were plated in poly-D-lysine coated 6-well plates at 400,000 cells/well in 2 mL/well to yield a confluency of 50% the following day. Media was changed to 1.5 mL/well DMEM containing 3% FBS and 1% NEAA. Cells were then transfected with 200 ng/well pCMV-VSVG, 800 ng/well pCMV-Gag/Pol, and 1000 ng/well pTRIP-CMV-Effector-IVSb-IRES-TagRFP using X-tremeGENE 9 (Sigma XTG9-RO) and OptiMEM (Thermo 31985062). Following 6 h of incubation, media was changed to 1.5 mL/well DMEM with 3% FBS and 1% NEAA. At 48 h post-media change, the supernatant was collected and replaced with fresh media. After an additional 24 h, the supernatant was again collected. Supernatants from both time points were pooled, centrifuged at 3000xg for 5 min to remove cell debris, and combined with HEPES to a concentration of 20 mM and polybrene to a concentration of 4 µg/mL. Lentivirus stocks were then arrayed in v-bottom 96-well plates and stored at -80°C.

### Statistical Analyses

Graphs were presented as mean values ± SEM with individual data points shown. At least three independent experiments were conducted for all quantitative experiments shown where statistical analyses were applied. All statistical analyses were performed with Prism software v10.0.2 (GraphPad) and statistical tests used are indicated in respective figure legends. Statistical significance (*P*<0.05) between compared groups is indicated in Figs as either: ns (not significant), *=P<0.05, **=*P*<0.01, ***=*P*<0.001, ****=*P*<0.0001.

## Supporting information

Table S1. Bacterial effector library screening results and cell viability data

Table S2. Key resources and reagents

## Acknowledgements

We thank Dr. Robert Orchard (UTSW Medical Center) for providing the pLentiCRISPRV2 vector. We thank Drs. Hanspeter Niederstrasser and Bruce Posner of the UTSW Medical Center High-Throughput Screening Core for assistance with experimental design and fluorescence imaging. We also thank Dr. Dahee Seo (UTSW Medical Center) for technical assistance. This work was supported by grants from the NIH (1R35GM137978 and 1R21AI169558) to DBG and by funding to DBG from the UTSW Endowed Scholars Program. NMA was supported by grants from the NIH (R01AI083359), Welch Foundation (I-1704), and Burroughs Wellcome Fund (1011019). AE and EAR were supported by NIH Training grant T32 AI007520. This work was also supported by grants to the UTSW Medical Center High-Throughput Screening Core (1S10OD018005-01 and 2P30CA142543-11).

## Author Contributions

**Aaron Embry**: Conceptualization; Data Curation; Formal Analysis; Funding Acquisition; Investigation; Methodology; Validation; Visualization; Writing – Original Draft; Writing – Review & Editing. **David Schad**: Data Curation; Formal Analysis; Investigation; Methodology; Writing – Review & Editing. **Emily A. Rex**: Data Curation; Formal Analysis; Funding Acquisition; Investigation; Methodology; Validation; Writing – Review & Editing. **Neal M. Alto**: Conceptualization; Formal Analysis; Supervision; Funding Acquisition; Methodology; Project Administration; Visualization; Writing – Review & Editing. **Don B. Gammon**: Conceptualization; Data Curation; Validation; Formal Analysis; Supervision; Funding Acquisition; Investigation; Methodology; Project Administration; Visualization; Writing – Original Draft; Writing – Review & Editing.

## Data Availability

All relevant data are within the manuscript or supporting data.

## Supporting Information

**Table S1. Bacterial effector library screening results and cell viability data.**

**Table S2. Key resources and reagents.**

### Supplementary Fig Legends

**Fig S1.**
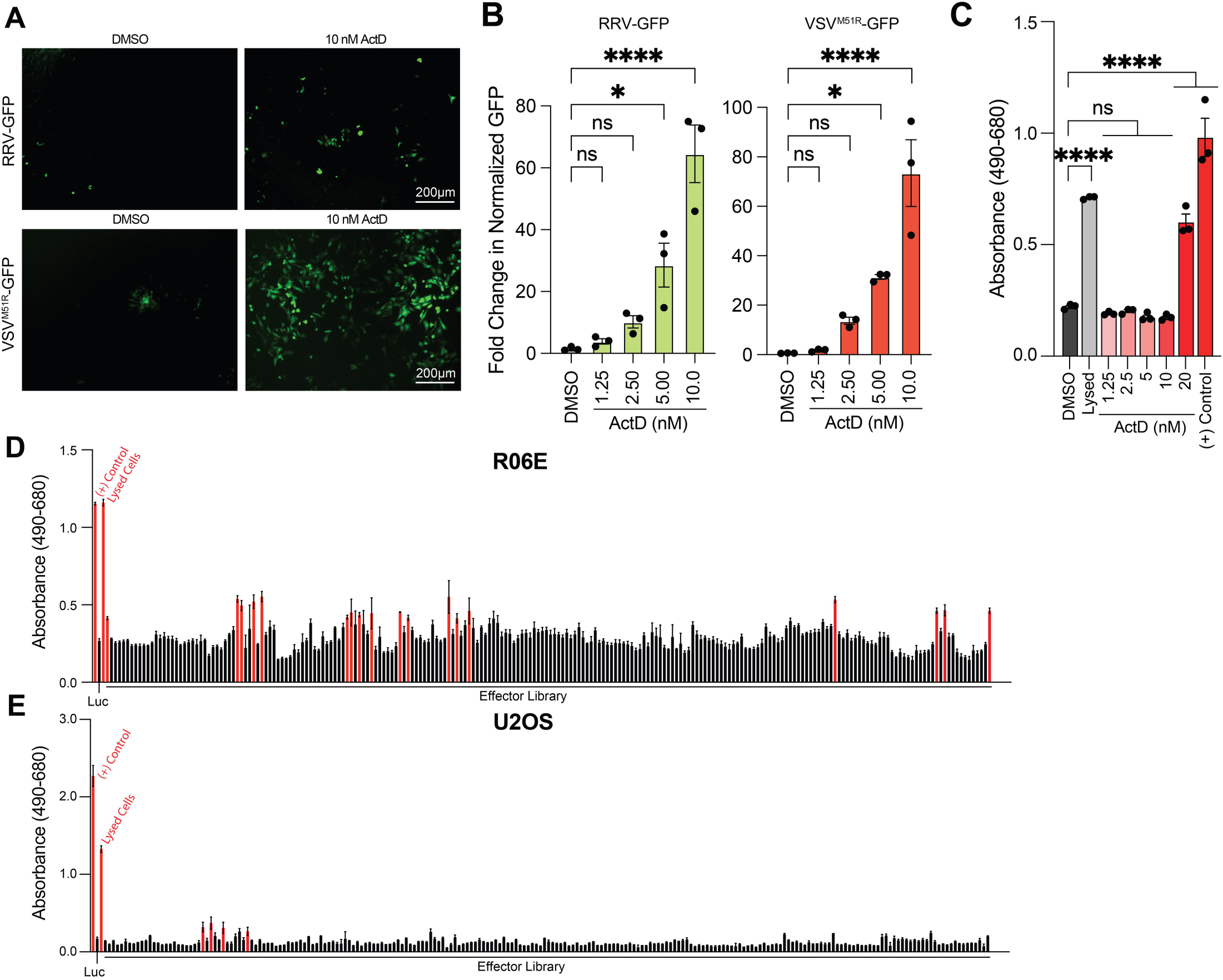
Effect of ActD treatment on arbovirus infection of mammalian cells and impact of effector expression on mammalian cell viability. **A.** Representative fluorescence microscopy images (GFP channel) of R06E cells infected with RRV-GFP (MOI=0.001) or VSV^M51R^-GFP (MOI=0.0001) and treated with either DMSO or 10 nM ActD for 20 h. Scalebars indicate 200 μm. **B.** Fold-change in GFP reporter readout following 20 h infection in the presence of the indicated doses of ActD, normalized to DMSO control. **C.** Results of LDH-based cytotoxicity assays in R06E cells. Absorbance at 490 nm is plotted for supernatant collected from R06E cells treated for 20 h with increasing doses of ActD. Positive (+) control supplied by the manufacturer, as well as cells lysed with manufacturer 10X lysis buffer (lysed cells) are also plotted for reference. **D.** Results of LDH-based cytotoxicity assays in R06E cells transduced with effector library for 48 h. LDH values that were significantly higher (P<0.05) from Luc-transduced control cells were considered toxic effectors (red bars). **E.** Results of LDH-based cytotoxicity assays in U2OS cells transduced with effector library for 48 h as in D. Red bars indicate toxic effectors. Data are means ± SEM; n=3. Statistical significance for B-C was determined with One-way ANOVA; ns (not significant), *=P<0.05, **=*P*<0.01, ***=*P*<0.001, ****=*P*<0.0001.

**Fig S2.**
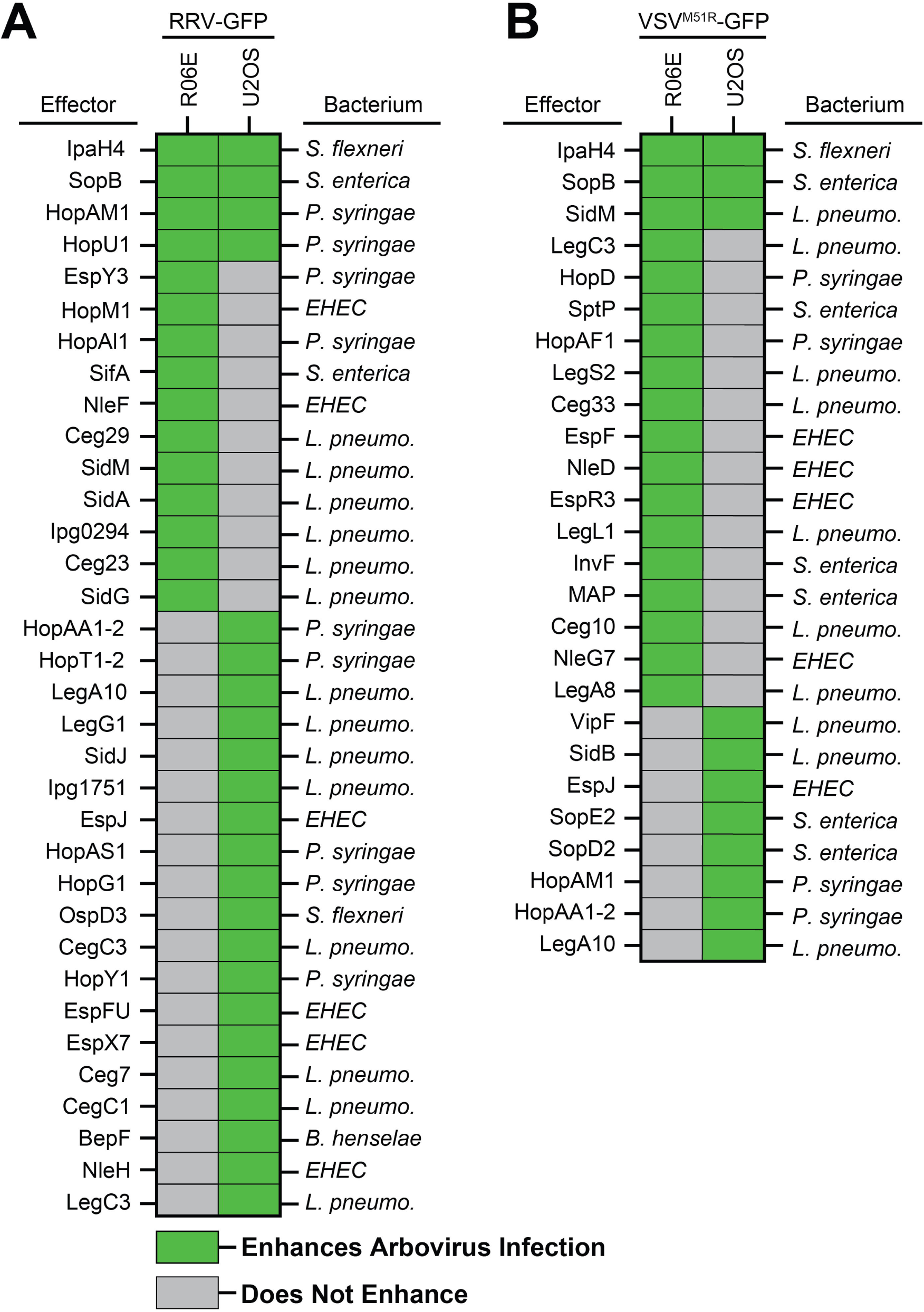
Specific bacterial effectors enhance arbovirus replication in bat and human cells. **A.** Summary of bacterial effectors that rescued RRV-GFP replication in bat or human cells, or both cell types. Green blocks indicate the effector enhanced RRV-GFP in the cell line indicated in the column header. Effectors are listed from high-to-low based on their fold-change in GFP signal over controls. The bacterium encoding each effector is noted to the right: *Shigella flexneri* (*S. flexneri*), *Pseudomonas syringae* (*P. syringae*), *Salmonella enterica* (*S. enterica*), *Legionella pneumophila* (*L. pneumo*.) *Enterohemorrhagic Escherichia coli* 0157:H7 (*EHEC*), or *Bartonella henselae* (*B. henselae*). **B.** Summary of bacterial effector proteins that enhanced VSV^M51R^-GFP replication in bat or human cells, or both cell types. The complete list of effectors screened and the raw results of the screens can be found in **Table S1**.

**Fig S3.**
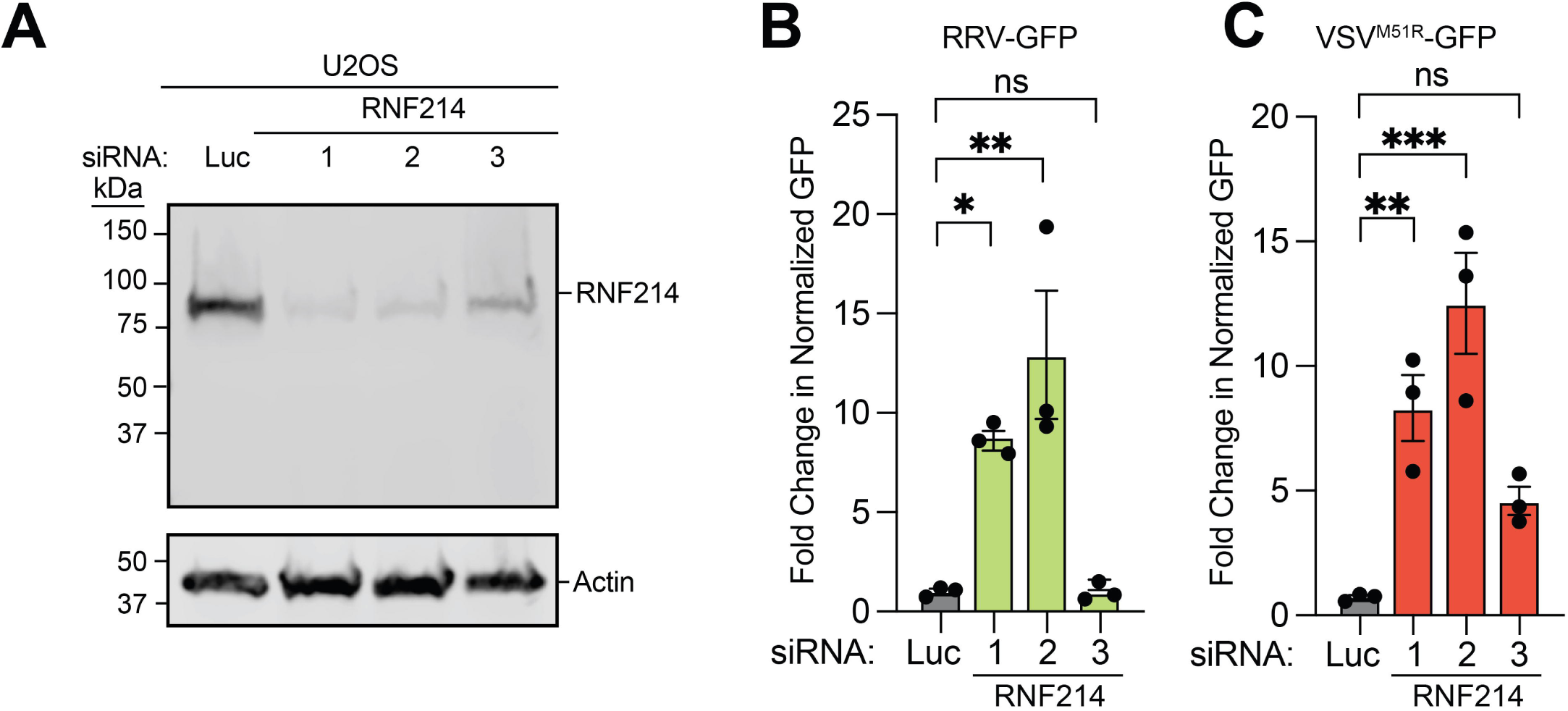
RNF214 knockdown enhances arbovirus replication in human U2OS cells. **A.** Representative immunoblot of RNF214 levels in U2OS cells transfected with indicated siRNAs for 48 h. **B.** Fold-change in GFP reporter readout following RRV-GFP infection of U2OS cells after indicated knockdowns. Cells were infected (MOI=0.001) 48 h post-siRNA transfection for 20 h, stained with CellTracker Orange, and imaged to determine the fold-change in GFP signal compared to Luc siRNA (control) treatments. **C.** Fold-change in GFP reporter readout following VSV^M51R^-GFP infection of U2OS cells after indicated knockdowns. Cells were infected (MOI=0.0001) 48 h post-siRNA transfection for 20 h, stained with CellTracker Orange, and imaged to determine the fold-change in GFP signal compared to Luc siRNA (control) treatments. Data are means ± SEM; n=3. Statistical significance was determined with One-way ANOVA; ns (not significant), *=P<0.05, **=*P*<0.01, ***=*P*<0.001, ****=*P*<0.0001.

**Fig S4.**
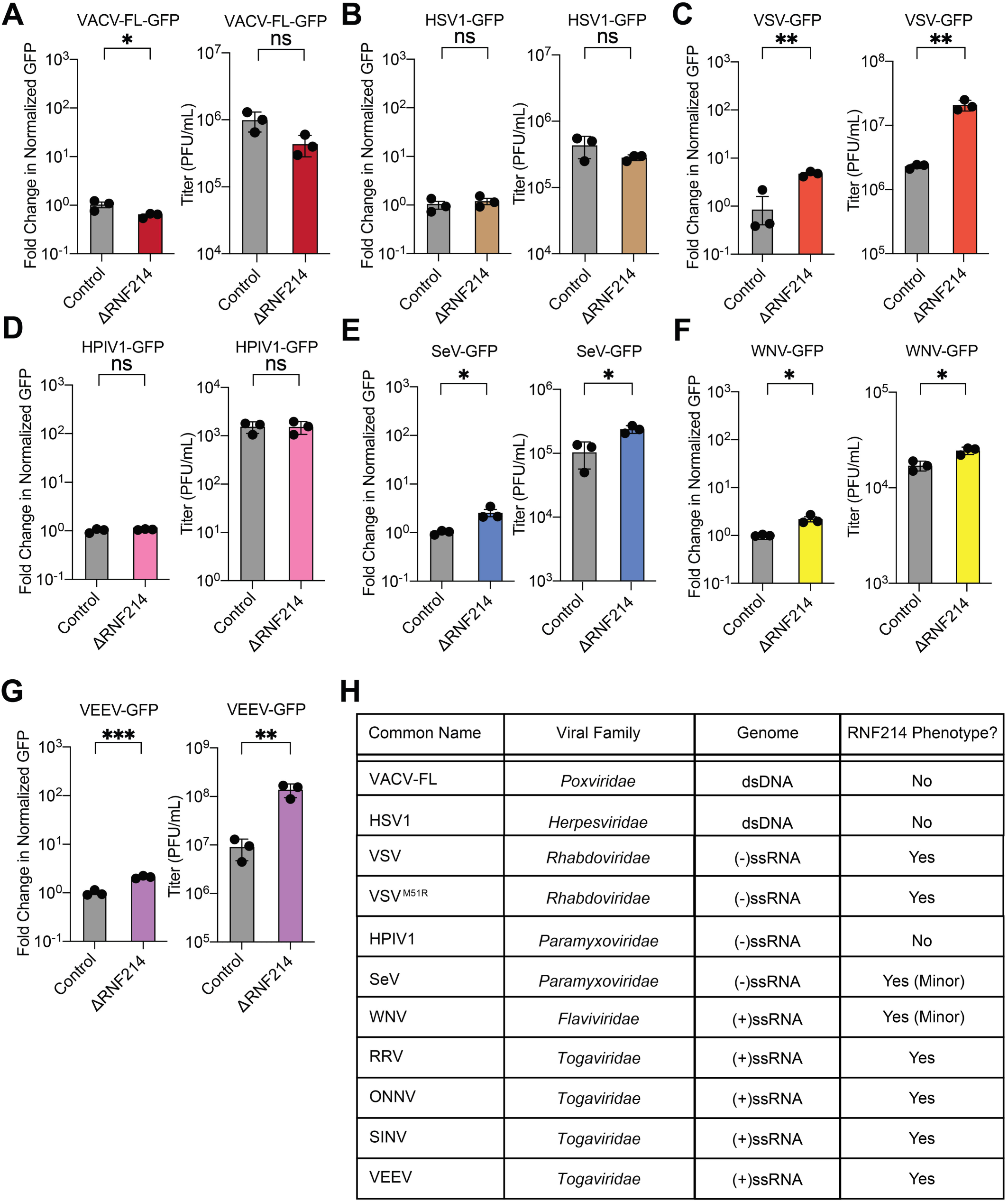
Assessing the impact of RNF214 knockout on the replication of diverse viral families. A-B. Control or U2OS^ΔRNF214^ cells infected with the indicated reporter poxvirus (VACV-FL-GFP; MOI=0.01) or herpesvirus (HSV-1-GFP; MOI=0.01). Following infection, cells were stained with CellTracker Orange and imaged for fold-change in GFP reporter readout compared to control cells. Titers are also shown from these collected cultures. **C-G.** Similar experiments were performed as in A-B but with VSV-GFP (MOI=0.0001) (C), the paramyxoviruses: HPIV1-GFP (MOI=0.001) (D) and SeV-GFP (MOI=0.001) (E), the flavivirus, WNV-GFP (MOI=0.001) (F) and the togavirus, VEEV-GFP (MOI=0.001) (**G**). **H.** Table summarizing results for the various viral families tested in this figure. Data for A-G are means ± SEM; n=3. Statistical significance was determined with unpaired student’s t-test; ns (not significant), *=P<0.05, **=*P*<0.01, ***=*P*<0.001, ****=*P*<0.0001.

